# Cooperativity in RNA chemical probing experiments modulates RNA 2D structure

**DOI:** 10.1101/2023.12.06.570415

**Authors:** Ethan B Arnold, Daniel D Cohn, Emma M Bose, Gregory Wolfe, Alisha N Jones

**Affiliations:** Department of Chemistry New York University 31 Washington Place, New York, New York 10003; Department of Physics New York University 726 Broadway, New York, New York 10003

**Keywords:** RNA structure, chemical probing, cooperativity, NMR, SHAPE, DMS

## Abstract

Small molecule chemical probes that covalently bond atoms of flexible nucleotides are widely employed in RNA structure determination. Atomistic molecular dynamic (MD) simulations recently revealed that the binding of RNA by chemical probes is influenced by cooperative effects, leading to measured reactivities that depend on the concentration of the chemical probe. In this work, we used selective 2’ hydroxyl acylation analyzed by primer extension (SHAPE) and dimethyl sulfate (DMS) chemical probing experiments to explore whether RNA structures are modulated by chemical probe binding events. We find that as the concentration of a chemical probe increases, modified nucleotides locally modulate the RNA structure, resulting in the increase or decrease of chemical probe reactivity in surrounding nucleotides. This cooperative effect is dependent on both chemical probe concentration and size. We find that these cooperative effects can be used to link structurally related nucleotides, and that the cooperative effects result in strikingly different 2D structure predictions as probe concentrations are varied.

## Introduction

Ribonucleic acid (RNA) has many regulatory roles in the cell that extend beyond protein translation; it also controls processes such as gene expression, chromatin remodeling, epigenetics, and RNA processing.^[1–3]^ It is through their primary sequences and secondary (2D) and tertiary (3D) structures that RNA interacts with proteins, other nucleic acids, and ligands to facilitate function.^[4,5]^ Thus, elucidating the structures of RNA not only provides insight into their function but guides the development of therapeutic agents when RNAs are implicated in disease.

Chemical probing has gained considerable traction as a means for determining the 2D structures and 3D interactions of an RNA.^[6–9]^ In these experiments, small molecule probes that target flexible and accessible nucleotides of an RNA in solution are used to obtain experimental restraints to guide the *in silico* prediction of RNA 2D structure.^[10–13]^ Briefly, an RNA in solution (or in cells) is treated with molecules that target specific atoms of flexible nucleotides. Dimethyl sulfate (DMS), for example, methylates the N1 and N3 atoms of adenosine and cytosine, respectively (**Scheme 1A**).^[14,15]^ Another class of probes acylates the 2’ hydroxyl group of flexible nucleotides through a technique known as selective 2’ hydroxyl acylation analyzed by primer extension (SHAPE) (**Scheme 1B**).^[16–20]^

**Scheme 1.**
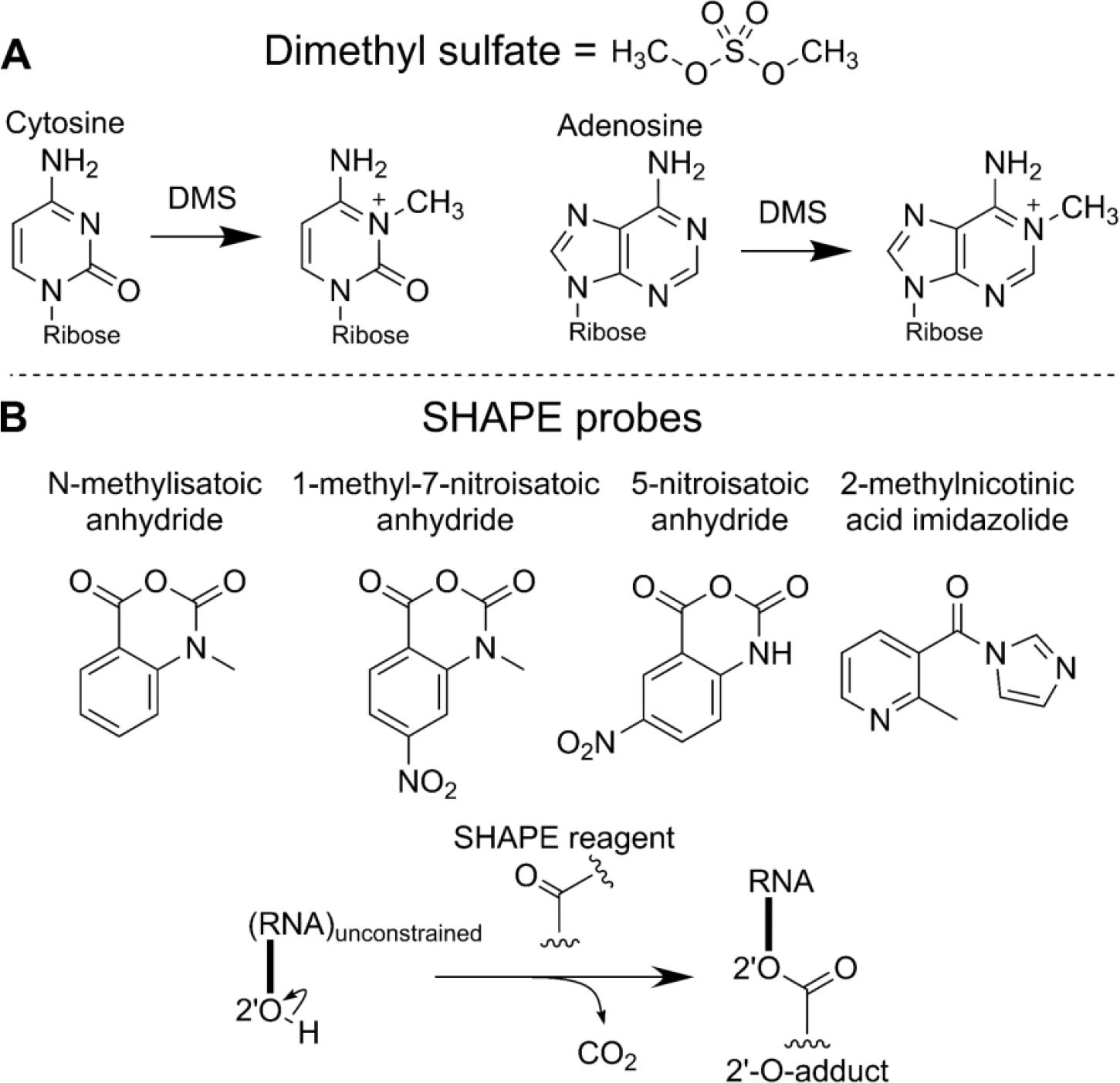
Mechanisms of RNA chemical probe modification. A) Dimethyl sulfate methylates the N1 and N3 atoms of flexible adenosine and cytosine residues, respectively. B) SHAPE probes acylate the 2’OH ribose group of flexible nucleotides.

Modified RNAs are then reverse-transcribed to reveal sites of modified nucleotides. Here, the reverse transcriptase will pause at modification sites (RT-stop) or introduce a mutation (RT-MaP).^[21]^ cDNA fragments resulting from paused elongation can be assessed via gel or capillary electrophoresis, and mutations are detected through next-generation sequencing approaches.^[17,22,23]^ The increased number of cDNA fragments or the high frequency of mutations at specific nucleotide positions generally correspond to a highly reactive (single-stranded or flexible) nucleotide. This information is then translated into a reactivity profile that serves as a guide for the *in silico* prediction of the RNA’s 2D structure.

In a recent computational study, the effect of 1-methyl-7-nitroisatoic anhydride (1M7) chemical probe binding on surrounding nucleotides was assessed.^[17,24]^ As the concentration of 1M7 increased, cooperative effects were observed; that is, the binding interaction of a nucleotide ‘i’ with 1M7 influenced the binding interaction of 1M7 on a neighboring nucleotide, ‘j’. In a non-cooperative scenario, a linear trend between probe concentration and reactivity is expected. A non-linear trend indicates a cooperative event.

Cooperativity is likely attributed to the distortion of structure imposed upon the RNA by both the concentration and bulkiness of the probe. Thus, in an ideal chemical probing experiment, low concentrations of probes should be used to ensure that single-hit kinetic modifications occur (where one modification occurs per molecule).^[25]^ This ideal concentration was demonstrated to be 1 mM or below for GNRA tetraloop RNAs based on molecular dynamics (MD) simulations.^[24]^ At non-ideal concentrations, the reactivity profile reflects a damaged RNA and the 2D structure of the RNA can be mispredicted.

## Methods

### In vitro transcription and purification of RNA

DNA templates encoding for the RNAs containing the T7 promoter region and 3’ and 5’ SHAPE cassettes were ordered from Integrated DNA Technologies (IDT).^[6]^ Sequences corresponding to the DNA templates are listed in Table 1.

**Table 1.**
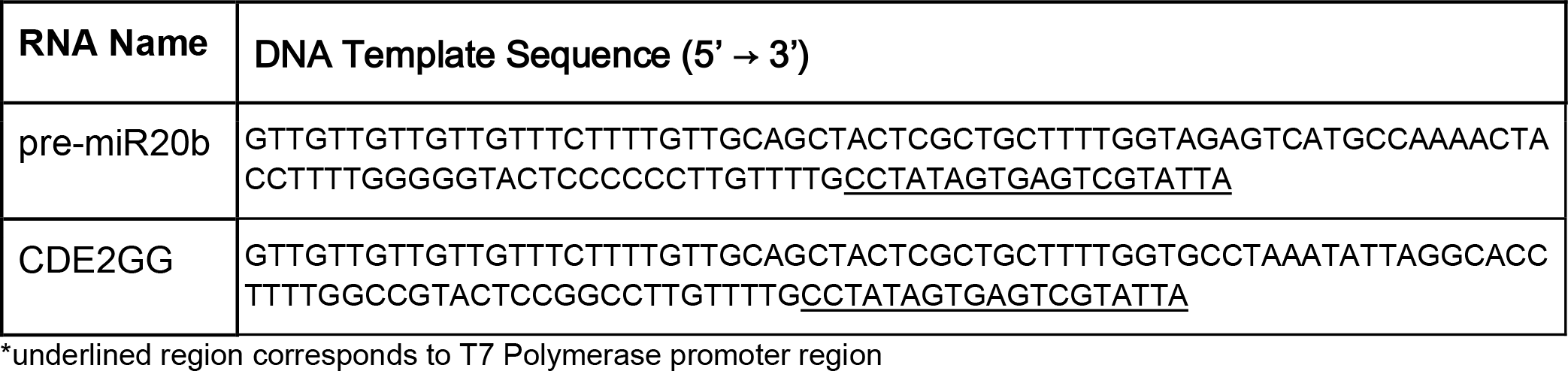
DNA templates purchased from IDT.

### Polymerase Chain Reaction using PRIMERIZE ^[26,27]^

For each 50 μL reaction, 100 μM of the outer forward (1F) and reverse (4R/8R) primers were combined with 1 μM of inner primers (2R-3F; 2R-7F) for a final concentration of 4 μM and 0.04 μM respectively, along with 5X Phusion Buffer (1X), 5X GC Enhancer (1X), 0.2mM dNTPs, and 0.5 μL Phusion Polymerase. Thermocycler settings were as followed: 1) initial denature at 98ºC for 30 sec, 2) denature at 98ºC for 10 sec, 3) anneal at 60ºC for 30 sec, 4) extension at 72ºC for 30 sec, 5) repeat steps 2-4 for 30 cycles, 6) final extension at 72ºC for 10 min. DNA products were checked on a 1% agarose gel, made in 1X TAE (tris(hydroxymethyl)aminomethane, acetic acid, disodium EDTA dihydrate) stained with SYBR Safe, and ran at 100V for 50 min.

RNAs were transcribed *in vitro* with bacteriophage T7 RNA polymerase (a gift from Michael Sattler). Briefly, the optimal MgCl_2_ concentration for transcription was determined by supplementing 0.64 μM of DNA template with 0.64 μM T7 promoter primer (5’s- TAATACGACTCACTATAGGG- 3’ ordered from IDT), 8 mM each rATP, rCTP, rGTP, rUTP, 5% polyethylene glycol 8000, 1X transcription buffer (5 mM Tris pH 8, 5 mM spermidine, 10 mM DTT), 20 to 80 mM MgCl_2_, 0.6 mg of T7 polymerase. For RNAs made through polymerase chain reaction (PCR), FIRRE RRD, T7 promoter primer was omitted. Following a 1 hr incubation at 37°C, transcriptions were purified on 8% polyacrylamide gels containing 8 M urea. RNA transcriptions yielding the most RNA were scaled up to produce RNA for chemical probing experiments. RNA was purified by denaturing PAGE, and bands containing RNA were excised from the gel and electro-eluted in 1X TBE buffer (100 mM Tris base, 100 mM boric acid, 2 mM EDTA). Ethanol precipitation was performed (to desalt the RNA) by adding 0.1 vol of 3 M sodium acetate, 3.3 vol pure ethanol, cooling at -80°C for 30 min, centrifuging for 1 hr at 4000 RPM at 4°C, decanting and aspirating the supernatant, and redissolving the RNA pellet in nuclease-free water. RNAs were stored at -20°C.

### Chemical probing experiments

RNAs were prepared by combining 1 μM RNA, H_2_O, folding buffer (100 mM NaCl, 50 mM HEPES pH 8, 0.1 mM EDTA ph 8, in H_2_O), snap cooling (3 min at 95ºC, 5 min on ice), adding refolding buffer (30 mM NaCl, 15 mM HEPES pH 8, 5 mM MgCl_2_, in H_2_O), and incubating at 37°C for 30 min. Chemical probes were diluted (SHAPE in dimethyl sulfoxide, DMS in 10% ethanol) to 15 mM, 25 mM, 50 mM, 75 mM, 100 mM, and 125 mM. 0.222 μM of prepared RNA mixture was treated with probe dilutions to yield a concentration of 0.2 μM RNA. Control reactions were carried out by replacing chemical probe with corresponding solvent. This was then incubated at 37°C, quenched by addition of 0.5 M Na-MES pH 6, 5 M NaCl, Invitrogen™ Poly(A)Purist™ Magnetic Beads, 0.25 μM fluorescein amidite primer, and H_2_O, incubated at room temperature for 10 min, then placed on a magnetic stand for 10 min. Supernatants were aspirated, and the pellets were washed twice with 70% ethanol. Air-dried pellets were resuspended in H_2_O.

Treated RNAs were reverse transcribed with Superscript III reverse transcriptase (SSIII, purchased from ThermoFisher) by addition of 5X reverse transcription buffer (250 mM Tris-HCl pH 8.3, 375 mM KCl, 15 mM MgCl_2_), 0.1 M dithiothreitol, 10 mM dNTP mix, SSIII reverse transcriptase (200 units/μL), H_2_O, followed by a 30 min incubation at 48°C. Reaction products were treated with 0.4 M NaOH, incubated at 95°C for 3 min, cooled on ice for 5 min, and quenched with a mixture of 1 vol 5 M NaCl, 1 vol 2 M HCl, and 1.5 vol 3 M sodium acetate. After a 7 min incubation at room temperature on a magnetic stand, supernatants were aspirated. Resulting cDNA was precipitated with ethanol, air-dried, resuspended in a mixture of 1000 parts formamide and 1 part 350 ROX dye standard, incubated at room temperature for 10 min, and placed on a magnetic stand for 7 min. Supernatant aliquots were placed in a new well plate, from which 3:20 dilutions were made in a mixture of 1000 parts formamide and 1 part 350 ROX dye size standard. Resulting saturated and diluted samples were analyzed by capillary electrophoresis.

Chemical probing experiments that were analyzed using mutational profiling were carried out as described in “Selective 2’-hydroxyl acylation analyzed by primer extension and mutational profiling (SHAPE-MaP) for direct, versatile and accurate RNA structure analysis.”^[28]^ Reverse transcription products were purified using RNAClean XP beads (Beckman Coulter), following the manufacturer’s protocol. Library preparation was done following the Small RNA Library Preparation, and each PCR product was purified using Ampure Beads (Beckman Coulter), following the manufacturer’s protocol. Integrity of the final PCR product was checked using the TapeStation and Quibit fluorometer. Sequencing was carried out on a MiSeq.

### Analysis of chemical probing data

HiTRACE was used to obtain chemical probe reactivities.^[29]^ Reactivity data was normalized by determining the first and third quartile and establishing the interquartile range (IQR). The upper extreme (UP) is then determined, and nucleotides with reactivities greater than UP are omitted. Following the removal of outliers maximum reactivity is commuted: the top 8% of reactivity values are averaged. Absolute reactivity values are divided by the maximum reactivity value to give normalized reactivity values at each nucleotide. Reactivity data can be found at Zenodo under the accession code: 10.5281/zenodo.10075871

### NMR Spectroscopy

CLEANEX-PM experiments were conducted using 100 μM unlabeled RNA to determine proton exchange rates (Kex). ^[30]^ Mixing times ranged from 5 to 175 ms, with the relaxation parameter held constant at 1 sec. Spectra were processed and analyzed using Topspin (Bruker), and peaks were picked and quantified using the equation illustrated below.^[30]^ R_1, A_ is the longitudinal relaxation rate for RNA, R_1,W_ is the longitudinal relaxation rate for water, and T_M_ is the mixing time. 2D ^1^H-^1^H NOESY and ^1^H, ^15^N HSQC experiments at 298K were conducted to assign imino proton resonance. CLEANEX data was fit in OriginPro graphing Software by the following equation:

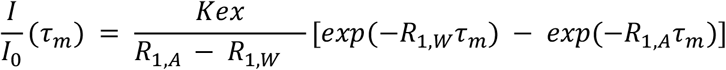

## Results

In this study, we experimentally investigated the impact of cooperativity as a function of chemical probe concentration for five chemical probes on the structures of three RNAs: the precursor microRNA (miRNA) 20b stem-loop (pre-miR20b), the constitutive decay element 2GG (CDE2GG), and the repeating RNA domain (RRD) of the functional intergenic repeating RNA element (FIRRE) long noncoding RNA (lncRNA) (**Table 3**). These chemical probes include DMS and four SHAPE probes: 1M7, N-methylisatoic anhydride (NMIA), 2-methylnicotinic acid imidazolide (NAI), 5-nitroisatoic anhydride (5NIA), and their effects were observed using RT-stop and RT-MaP experiments.^[6,14,17,19,20,31]^

**Table 2.**
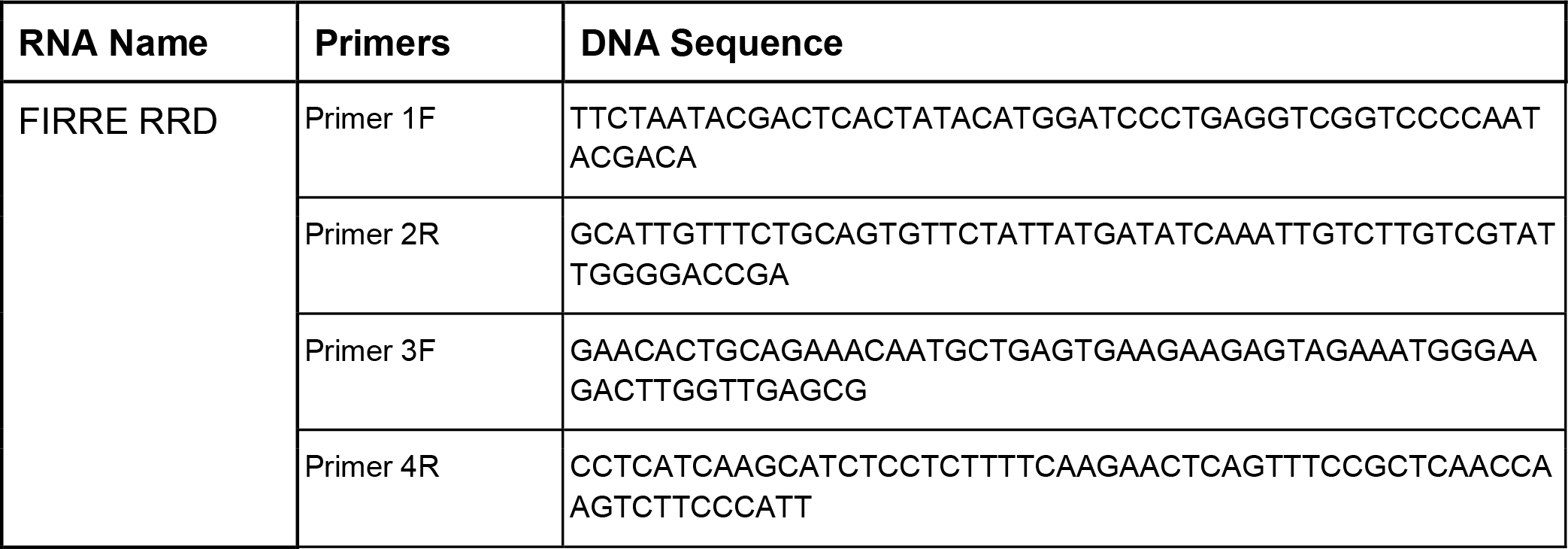
DNA primers purchased from IDT for Polymerase Chain Reaction (PCR)

**Table 3.** RNAs investigated in this study.

| RNA | Length |
| --- | --- |
| Precursor microRNA 20b stem-loop (pre-miR20b) | 23 nt |
| Constitutive Decay Element 2GG | 21 nt |
| Functional Intergenic Repeating RNA Element Repeating RNA Domain (RRD) | 154 nt |

Both the CDE2GG and pre-miR20b RNAs fold to form stem-loop structures.^[32,33]^ Pre-miR20b is responsible for miRNA biogenesis involved in tumor suppression; CDE2GG is recognized as a constitutive decay element (CDE) by RNA binding protein (RBP) Roquin mediating mRNA degradation for autoimmune responses.^[32,34,35]^ We carried out RT-stop chemical probing experiments at six different concentrations of probe (1.5, 2.5, 5.0, 7.5, 10.0, and 12.5 mM SHAPE and 1%, 0.94%, 0.88%, 0.82%, and 0.78% DMS) and determined the reactivity at each concentration (**Figure 1A-B**). The following trends were observed:

**Figure 1.**
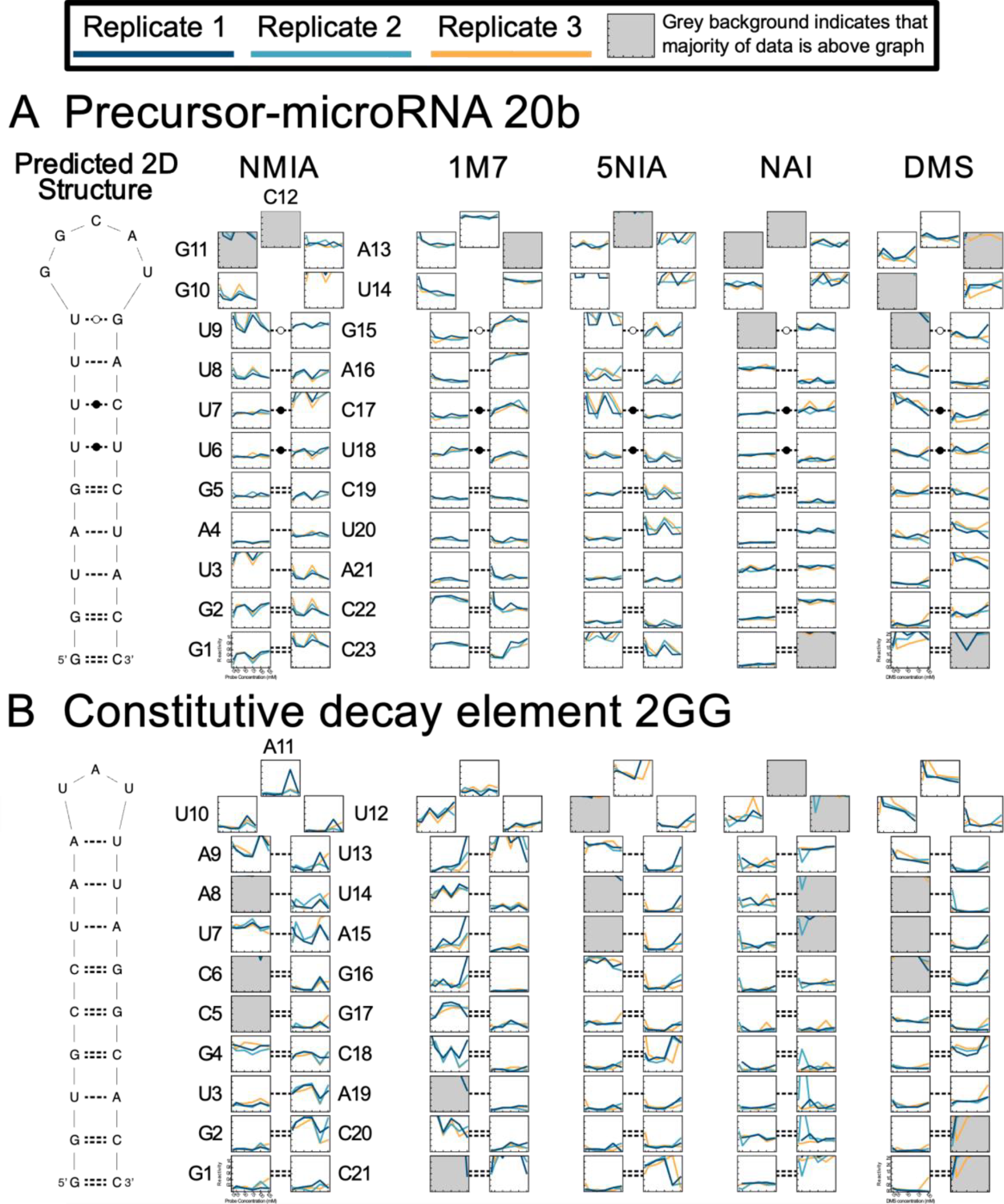
Reactivity of each nucleotide in A) pre-miR20b, B) CDE2GG as a function of chemical probe concentration for NMIA, 1M7, 5NIA, NAI, and DMS. Each small graph represents one nucleotide, and the graphs are arranged in the shape of their RNA’s hairpin.

*i) nucleotides in both the stem and apical loop are subject to cooperative and anti-cooperative events:* A non-linear trend in reactivity, as the concentration of probe increases, indicates a cooperative or anti-cooperative event. This was observed for several nucleotides, occurring in both base-paired and single-stranded regions. For example, we refer to base-paired residues U8 and A16 of pre-miR20b RNA, which show a nonlinear decrease and increase in 1M7 reactivity respectively. U8 reflects an anti-cooperative event, whereas A16 is cooperative. A nonlinear decrease in reactivity is observed for pre-miR20b G10, which is in the apical loop, indicating an anti-cooperative event.

*ii) nucleotides opposite each other showed opposite trends in reactivity*. In the lower stem of pre-miR20b, the G2-C22 base pair possesses opposite trends in reactivity as the concentration of the 1M7 probe increases. This may be caused by probes disrupting stacking interactions between nucleotides, rendering the proximal structure more flexible. Notably, this opposite trend may offer a means for identifying structurally related nucleotides.

*iii) non-cooperative effects were generally observed for NAI relative to other chemical probes*. This may be due to the stabilized structure of the chemical probe, which is different from the others (**Scheme 1B**). In a previous study, NAI was reported to possess an electron-deficient structure, thus increasing the electrophilicity of the carbonyl group.^[20]^

We found it interesting that several nucleotides in stem regions are reactive with the chemical probe, whereas others are not, even at low probe concentrations. This indicated that some nucleotides in the stem may be more intrinsically dynamic than others, enabling their acylation or methylation by the chemical probes. Using nuclear magnetic resonance (NMR) spectroscopy chemical exchange (CLEANEX) experiments, we assessed the flexibility of these base pairs (**Figure 2**). CLEANEX experiments revealed that base-paired nucleotides that exhibit chemical probing reactivity are more flexible in solution; their iminos have a high rate of exchange with water protons. These results explain a frequently observed effect in chemical probing experiments, wherein base-paired nucleotides unexpectedly exhibit increased reactivity.

**Figure 2.**
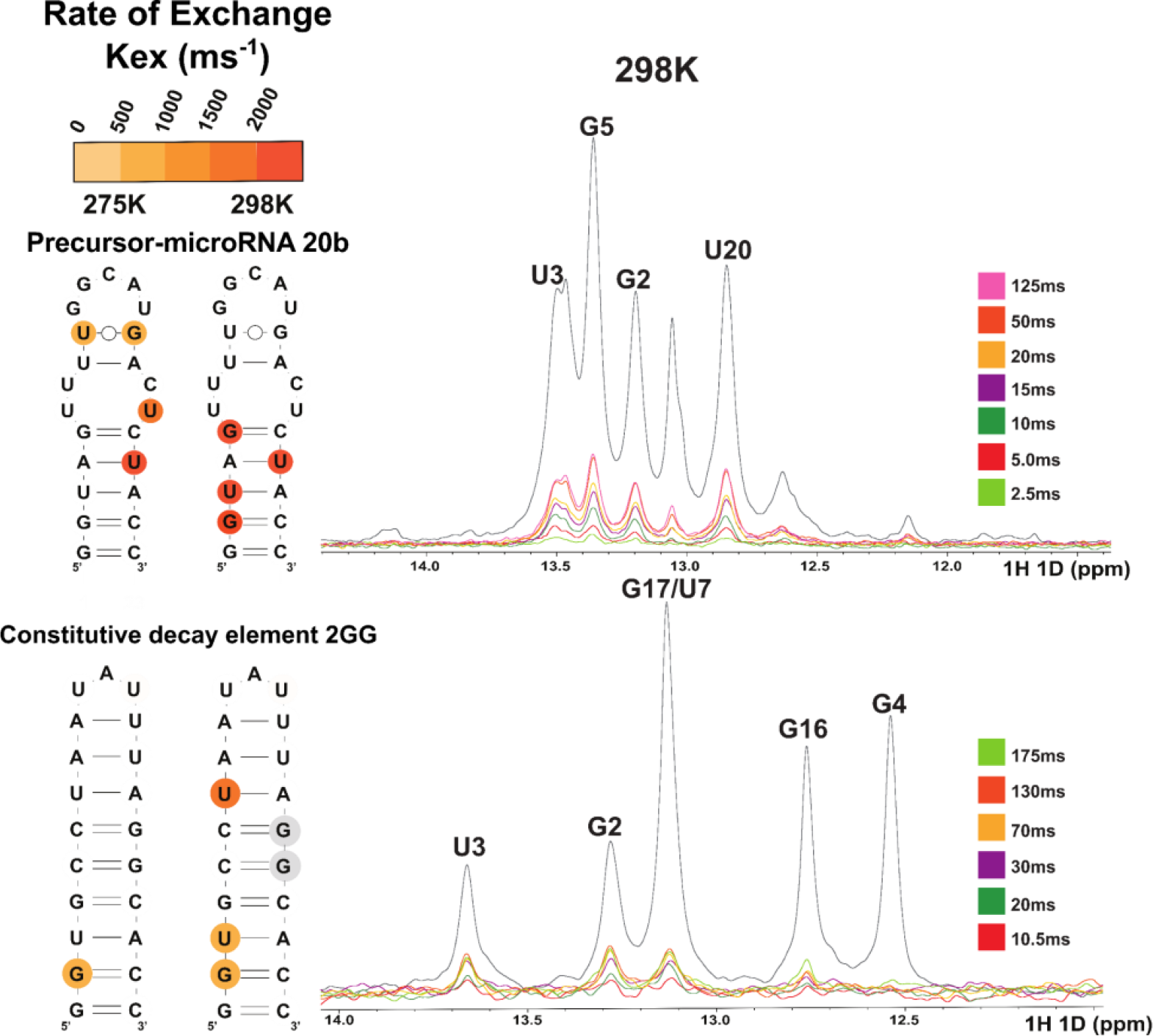
CLEANEX experiments for pre-miR20b (top) and CDE2GG (bottom). Experiments were conducted at 275K and 298K, with the D8 parameter ranging from 5 ms to 175 ms, to determine which imino resonances show higher rates of exchange (Guanosines and Uracils). 1D spectra shown for experiments ran at 298K. The 2D structures for each RNA are shown, as well as the nucleotide’s rate of exchange (Kex) with water highlighted in different shades of orange.

A noteworthy observation of the probing results on the stem-loop structured RNAs is the shift of a nucleotide’s reactivity, from non-reactive to reactive or the reverse, when probe concentration is varied (**Figure 3A**). Effectively, the reactivity profiles are different for RNAs at low concentrations of the probe relative to higher concentrations of the probe, which results in different 2D structure predictions. Indeed, the lowest energy 2D structure of the stem-loop of pre-miR20b is predicted to fold into different structured states as the concentration of the 1M7 chemical probe increases (**Figure 3B**). In particular, nucleotides associated with the upper region of the stem show differences in base pair patterns as the concentration of chemical probe increases.

**Figure 3.**
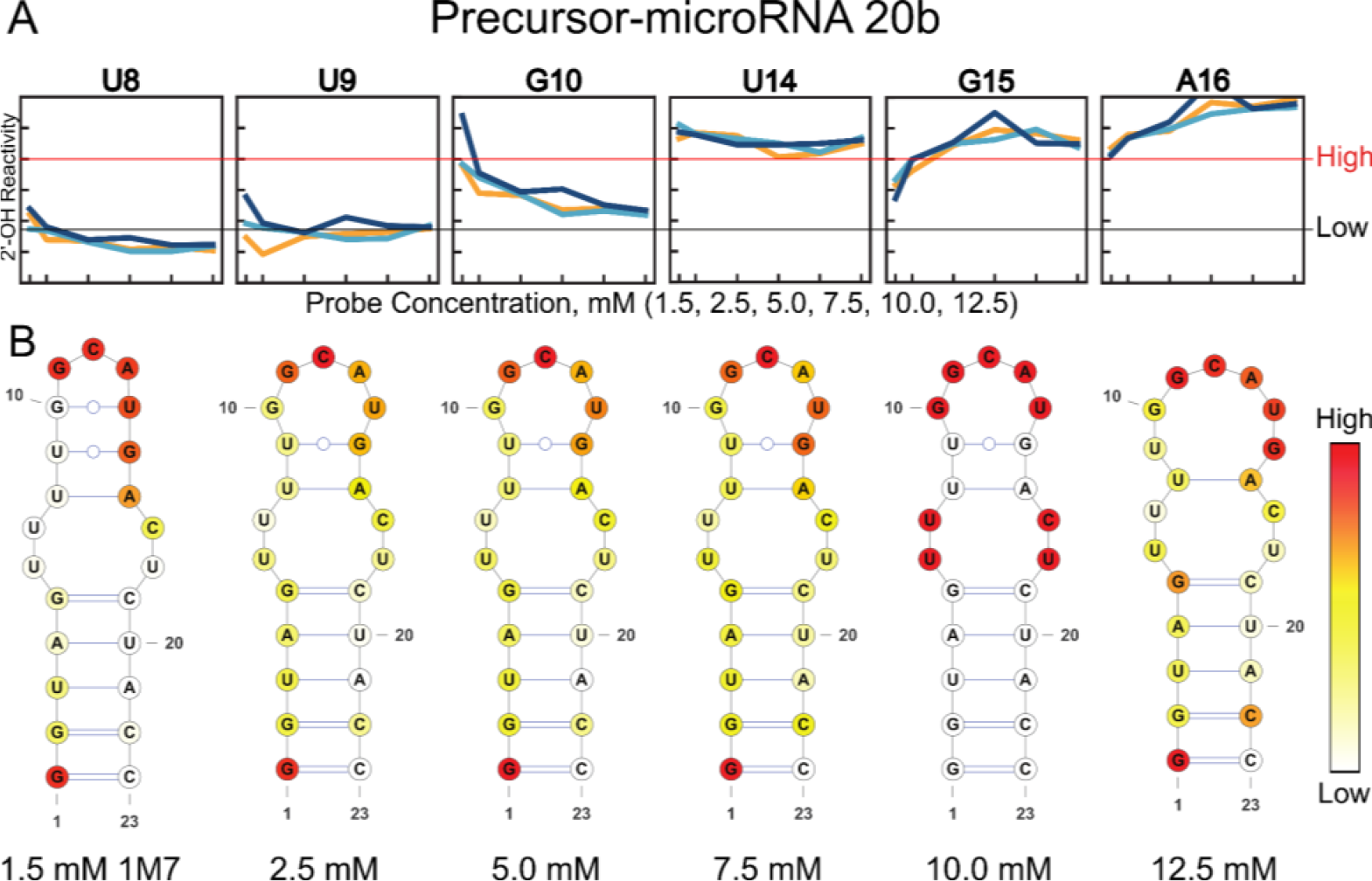
Impact of probe concentration on loop nucleotide reactivities and predicted 2D structure. A) Reactivity plots of nucleotides adjacent to the apical loop of pre-miR20b. “High” indicates reactivity >0.8; low indicates reactivity below <0.35. B) Lowest free energy predicted 2D structures of pre-mirR20b constrained by 1M7 SHAPE data.

This observed effect is not only seen in RT-stop experiments but also in RT-MaP. The FIRRE lncRNA is implicated in regulating several biological processes, including nuclear chromosome organization, adipogenesis, and hematopoiesis.^[36–39]^ Through a ∼156-nt domain known as the RRD, it interacts with several proteins to facilitate function.^[40]^ We increased the concentration of the 1M7 and DMS chemical probes and assessed the predicted structures: with increases in probe concentration, the reactivity profile is modulated, and the predicted structures in the ensemble vary (**Figure 4**). At the lowest concentration of 1M7, the FIRRE RRD is predicted to form a five-stem multi-way junction. However, as the concentration of 1M7 increases, the RRD folds into an elongated stem-loop structure, possessing several internal loops and bulges. The structured predicted based on DMS chemical probing are starkly different from 1M7, and also show variable structures as the DMS concentration is increased. In each of these structures, the RNA folds to form multiple branched helical domains. The differences in structure between the 1M7 and DMS-predicted structures may be attributed to the limited probing data; 1M7 provides information regarding all flexible nucleotides in solution, whereas DMS is limited to flexible adenosine and cytosine residues. Nevertheless, cooperative effects are observed for both DMS and SHAPE modifying probes in RT-MaP chemical probing experiments, and can change the outcome of structure predictions.

**Figure 4.**
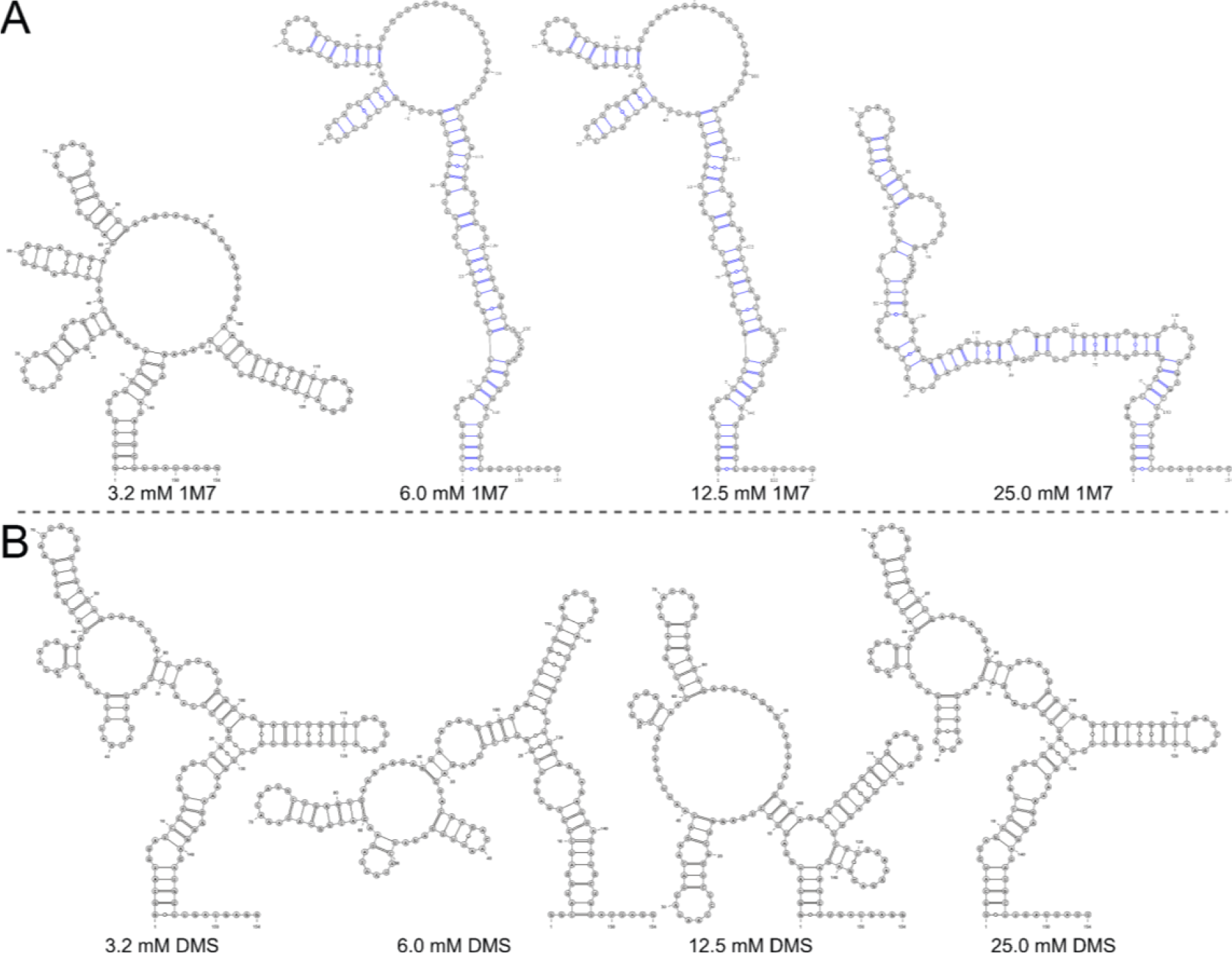
2D structure predictions differ between various chemical probe concentrations. The structures depicted are the free energy-predicted 2D structures of the FIRRE RRD probed with A) 1M7 and B) DMS.

## Discussion

This work highlights the impact of cooperativity on RNA structure prediction. In single-hit kinetic experiments, as the concentration of the chemical probe increases, a linear trend in reactivity is anticipated. However, we observe nonlinear trends for each of the chemical probes investigated in this study. Specifically, when applying our probing results as experimental restraints for thermodynamic folding algorithms, we detected divergences in the RNA structures as we varied probe concentrations. In a word, the reactivity profiles resulting from RNA chemical probing experiments are significantly impacted by probe concentration, due to cooperative effects confounded by each probe’s structure and identity.

Though chemical probing experiments are useful for gaining insight into the structure of RNA, it is important to maximize their accuracy. The concentration and type of a chemical probe during experimental design must be considered in order to mitigate cooperative effects. Several studies have been carried out to assess the bias of chemical probes for specific nucleotides, and programs (such as IPANEMAP) have been developed to account for the behaviors of probes in 2D structure prediction.^[19,41]^

The consequence of cooperativity is not just mispredicted 2D structures: many RNA 3D structure prediction programs rely on 2D structural data. Auto-DRRAFTER, for example, is used to build RNA coordinates into cryogenic electron microscopy (cryo-EM) maps.^[42]^ Incorrectly predicted 2D structures can have negative effects on 3D classification in cryo-EM analysis. Furthermore, incorrect structures detriment our understanding of how RNAs function through structure and our ability to carry out structure-based drug design. It is our recommendation that when chemical probing experiments are used to study RNA 2D structure, multiple probe concentrations should be assessed and reported.

## Supporting Information

The authors have cited additional references within the Supporting Information.^[6,29,30]^

## Acknowledgments

The authors would like to thank Katherine Buchan for assistance in collecting NMR spectra, the lab of Evgeny Nudler for assistance with MiSeq sequencing, Emma Gogarnoiu for assistance with schemes, and members of the Jones research group and the labs of Giovanni Bussi, Danny Incarnato, and Steve Bonilla for useful discussions.

## Notes

Supporting information for this article is given via a link at the end of the document.

https://zenodo.org/records/10075872

### Competing Interest Statement

The authors have declared no competing interest.

