## Supporting Information for "Cooperativity in RNA chemical probing experiments modulates RNA 2D structure"

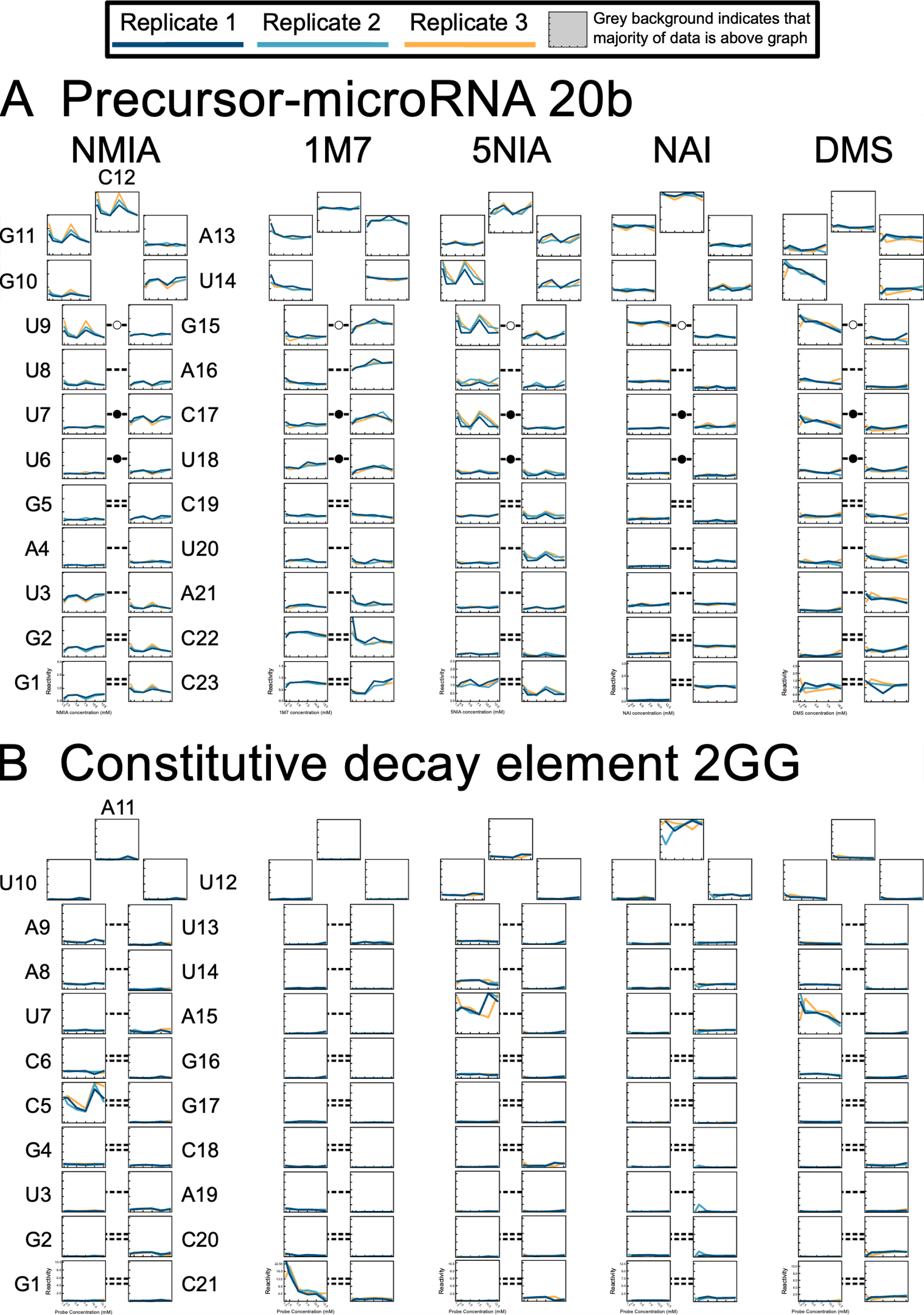

**Supporting Figure 1:** Chemical probe reactivity profiles for A) pre-miR20b and B) CDE2GG as functions of probe concentration for five chemical probes (NMIA, 1M7, 5NIA, NAI, and DMS); Graph bounds fit to data extremes.

**Supporting Figure 2:** CLEANEX experiments for pre-miR20b (top) and CDE2GG (bottom). Experiments were conducted at 275K and 298K, with the D8 parameter ranging from 5 ms to 175 ms, to determine which base-paired nucleotides (Guanosines and Uracils) show higher rates of exchange. The 2D structures for each RNA is shown, as well as the nucleotide’s rate of exchange (Kex) with water highlighted in different shades of orange.

**Supporting Figure 3:** Fitted curves done in OriginPro Graphing Software to determine the Rate of Exchange (Kex) for pre-miR20b (top) and CDE2GG (bottom) at two temperatures, A) 298K and B) 275K). Plots show the exponential growth of peak intensity as the mixing time (ms) increases from 5 ms to 175ms.
